# Entry, replication and innate immunity evasion of BANAL-236, a SARS-CoV-2-related bat virus, in *Rhinolophus* and human cells

**DOI:** 10.1101/2025.09.29.679146

**Authors:** Ségolène Gracias, Samuel Donaire-Carpio, Françoise Vuillier, Elodie Le Seac’h, Léa Vendramini, Adam Moundib, Magdalena Rutkowska, Anastasija Cupic, Javier Juste, Sarah Temmam, Flora Donati, Carles Martinez-Romero, Nathalie Morel, Olivier Schwartz, Nevan J Krogan, Lisa Miorin, Caroline Demeret, Philippe Roingeard, Sandie Munier, Jyoti Batra, Adolfo Garcia-Sastre, Vincent Caval, Nolwenn Jouvenet

## Abstract

Asian *Rhinolophus* bats are considered the natural reservoirs of an ancestral SARS-CoV-2. However, the biology of SARS-CoV-2-related viruses in bat cells is not well understood. Here, we investigated the replication of BANAL-236, the only bat-derived SARS-CoV-2 relative isolated to date, in *Rhinolophus* cells. BANAL-236 did not replicate in wild-type *Rhinolophus* cell lines. Entry assays using pseudoviruses expressing the spike proteins (S) of SARS-CoV-2, BANAL-236, and BANAL-52 revealed that efficient S-mediated entry depends on the expression of human ACE2 (hACE2) and human TMPRSS2 (hTMPRSS2) in human and *Rhinolophus* cells. Expression of *Rhinolophus* entry factors, either alone or in combination, did not facilitate SARS-CoV-2 or BANAL-236 entry in human cells, suggesting that the S protein of BANAL-236 interacts more efficiently with hACE2 than with its *Rhinolophus* counterpart (rACE2). Through biochemical, virological, and electron microscopy analyses, we showed that BANAL-236 and SARS-CoV-2 completed their replication cycles in a *Rhinolophus* cell line engineered to express high levels of hACE2 and hTMPRSS2. Despite efficient viral replication in modified *Rhinolophus* and human cells, no induction of interferon (IFN)-stimulated genes was detected. Using a screening approach, we identified several BANAL-236 proteins that antagonize IFN production and signaling in human cells. Our findings thus show that BANAL-236 possesses critical features that enabled zoonotic spillover: hACE2 usage and potent evasion of human IFN responses. The *Rhinolophus* cellular model we established offers a platform for further investigating the interactions between bat coronaviruses and their reservoir hosts.

**Author summary:** Bats are known reservoirs for viruses that cause severe diseases in humans, such as coronaviruses and filoviruses. Bat species naturally or experimentally infected with these viruses rarely exhibit clinical symptoms, suggesting an evolved tolerance to viral infections. To elucidate the mechanisms underlying viral tolerance and to identify factors that could facilitate zoonotic spillover, it is essential to study the replication of bat-borne viruses in relevant bat cellular models. Here, we investigated the replication of BANAL-236, a SARS-CoV-2 related virus isolated from fecal samples of *Rhinolophus* bats in Northen Laos, in a novel cell line derived from *Rhinolophus ferrumequinum* lung fibroblasts. Our findings reveal that BANAL-236 can efficiently use human entry factors and potently evade the human innate immune response, two traits that may have contributed to its zoonotic transmission. Furthermore, the *R. ferrumequinum* cell lines we developed is a valuable model for investigating the molecular interactions between sarbecoviruses and their natural hosts.

## Introduction

Severe acute respiratory syndrome coronavirus (SARS-CoV) and SARS-CoV-2 both belong to the subgenus *Sarbecoviruses* within the *Betacoronavirus* genus. They have the potential to cross species barriers, as demonstrated by their emergence in the human population in 2003 and 2019, respectively. In 2020, seven sarbecoviruses, collectively referred to as BANAL-CoVs, were recovered from fecal samples of *Rhinolophus* bats in Northern Laos [1]. Three of these viruses are the closest known relatives of the Wuhan strain of SARS-CoV-2: *R*. *malayanus* BANAL-52, *R*. *pusillus* BANAL-103 and *R*. *marshalli* BANAL-236 [1]. An ancestral SARS-CoV-2 may have originated from recombination events involving these viruses and other closely related *Rhinolophus* sarbecoviruses [1].

The spike (S) glycoprotein of sarbecoviruses mediates the attachment to and fusion with the host cell membrane. During its biosynthesis in the Golgi of infected cells, the S protein of SARS-CoV-2 is cleaved by proprotein convertases, such as furin, into S1 and S2 subunits [2], which remain non-covalently associated on the surface of mature virions. The S protein of BANAL-CoVs lack the furin-cleavage site (FCS) between the S1 and S2 subunits [1]. The receptor-binding domain (RBD) within the S1 subunit determines the binding affinity to the surface receptor angiotensin-converting enzyme 2 (ACE2) and is therefore a key determinant for host range and pathogenesis. Seventeen residues, so called ‘contact residues’ are essential for the RBD-ACE2 interaction [2]. The RBD of BANAL-236 differs from that of SARS-CoV-2 by only a single contact residue, yet it binds strongly to human ACE2 (hACE2) and facilitates hACE2-dependent entry and replication into human cells [1]. Binding of the S protein to ACE2 induces conformational changes in both subunits, exposing a S2 cleavage site called S2’. This cleavage is mediated by the transmembrane protease serine 2 (TMPRSS2) at the cell surface, triggering the formation of a fusion pore that releases into the cytosol the positive sense, single-stranded viral genome complexed with nucleocapsid (N) proteins [2,3]. In the absence of cell surface TMPRSS2, the virus enters cells via endocytosis and S2’ is cleaved by endosomal proteases, such as cathepsin L [2,3]. BANAL-236 and BANAL-52 exhibit a greater dependence on TMPRSS2 than SARS-CoV-2 in Vero cells expressing both ACE2 and TMPRSS2 [4].

BANAL-236 was successfully isolated by inoculating rectal swabs of *R*. *marshalli* on trypsin-treated Vero-E6 cells [1], which are African green monkey kidney cells widely used for viral propagation. Replication in these cells was dependent on ACE2 [1]. BANAL-236 also replicated in human intestinal Caco-2 and Calu-3 cells, both of which express endogenous ACE2 [1]. Studies using infectious molecular clones of BANAL-236 and BANAL-52 showed that both viruses replicated less efficiently than SARS-CoV-2 in primary human nasal epithelial cells and in the upper airway of transgenic mice expressing hACE2, as well as in both the upper and lower airways of Syrian golden hamsters [4]. Furthermore, BANAL-236 and BANAL-52 exhibited reduced pathogenicity and pneumotropism compared to SARS-CoV-2 in hACE2-expressing mice and Syrian golden hamsters [4–6] and showed limited transmission among Syrian golden hamsters [4]. The attenuated replication and transmission of BANAL-CoVs cannot be attributed to poor affinity for host receptors, as BANAL-236 S protein binds both hamster and human ACE2 with higher affinity than the S protein of early SARS-CoV-2 isolates [1,5].

Mammalian cells have evolved robust antiviral defense mechanisms, including the interferon (IFN) response, to detect and control viral infections. The recognition of viral nucleic acids by immune sensors initiates a signaling cascade that culminates in the production and secretion of IFNs. These secreted IFNs bind to receptors on the surface of both infected and neighboring cells, activating the JAK/STAT pathway. This, in turn, induces the expression of approximately 2000 IFN-stimulated genes (ISGs) [7]. Many ISGs act to disrupt the viral life cycle by targeting specific stages of replication, such as viral entry, protein translation, genome replication or assembly of new virions. High-throughput loss– and gain-of-function screens in human cells have identified several human ISGs with potent antiviral activities against SARS-CoV-2 [8]. For instance, LY6E inhibits S-mediated fusion of SARS-CoV-2 and other coronaviruses in both human cells and mice [9], while the prenylated form of OAS1 activates the endoribonuclease RNase-L to degrade SARS-CoV-2 RNA [10]. The coordinated action of ISGs efficiently controls SARS-CoV-2 replication. Accordingly, impaired IFN responses in COVID-19 patients are associated with an increased risk of severe disease [11].

Like all viruses, sarbecoviruses have evolved strategies to evade the IFN response in human cells [12–15]. A substantial proportion of SARS-CoV and SARS-CoV-2 proteins are capable of inhibiting key components of IFN pathways [12,16–20]. For example, the viral proteases NSP3 and NSP5 cleave IRF3, thereby suppressing IFN production [21–23]. As a result, SARS-CoV-2 is a weak inducer of the IFN response in some human cell models, such as hACE2-expressing A549 lung cells and human lung tissues, compared to other respiratory RNA viruses [24–26]. In contrast, SARS-CoV-2 replication in bat cells derived from *Eptesicus serotinus, Eptesicus fuscus* and *Myotis myotis* expressing hACE2 triggers ISG expression [25,27], indicating that viral IFN counteraction mechanisms may be species-specific. However, the ability of BANAL-CoVs to counteract the IFN response in *Rhinolophus* cells remains unexplored.

Here, we investigated the replication of BANAL-236 in cells isolated from *Rhinolophus* bats, with a focus on viral entry mechanisms and the evasion of innate immunity.

## Results

### BANAL-236 and SARS-CoV-2 failed to replicate in RFe cells

To better understand the biology of BANAL-236 in a relevant cellular model, we first established a primary lung fibroblast cultures from the lung tissues of *a R. ferrumequinum* (RFe) bat collected in Spain [28]. The cells were then immortalized by expressing SV40-LT antigen [28]. To assess the expression of key viral entry factors, we performed RT-qPCR analysis to quantify endogenous ACE2 and TMPRSS2 mRNA levels in RFe cells, as well as in 293T and Caco-2 cells as comparisons. Consistent with previous reports [29], 293T and Caco-2 cells expressed moderate levels of ACE2, approximately10^2^ copies per μg of total RNA (Fig. 1A), whereas ACE2 mRNA was undetectable in RFe cells. Both 293T and RFe cells expressed less than 10 copies of TMPRSS2 per μg of total RNA. By contrast, Caco-2 cells expressed high levels of the protease, over 10^3^ copies per μg of total RNA (Fig. 1A).

**Figure 1.**
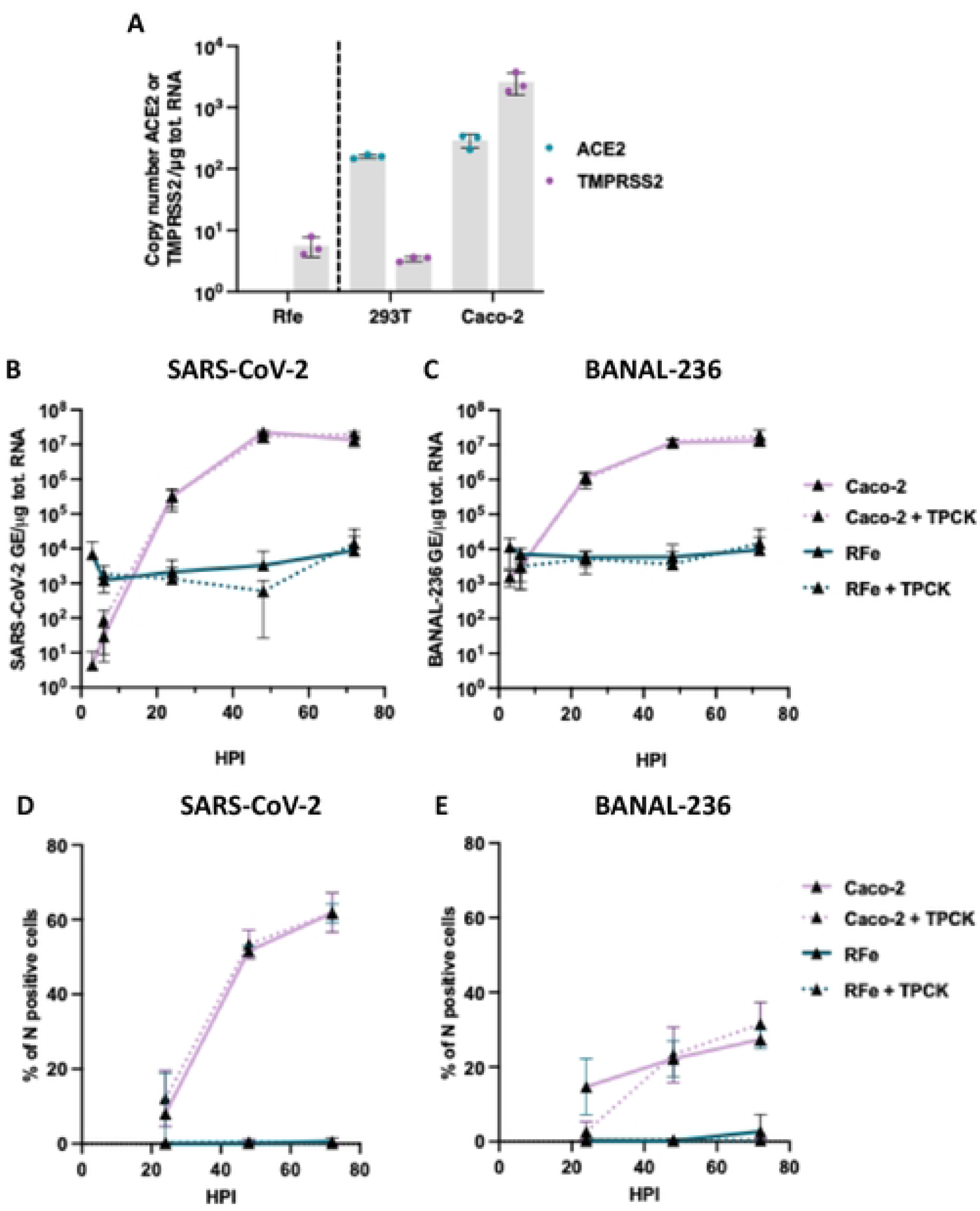
SARS-CoV-2 and BANAL-236 replicate in Caco-2 cells, but not in RFe cells. (**A**) Quantification of copy numbers per µg of total cellular RNA of endogenously expressed ACE2 and TMPRSS2 in indicated cell lines via RT-qPCR analysis. Data are means ± SD of three independent experiments. Caco-2 cells (pink lines) were infected with SARS-CoV-2 (**B-D**) or BANAL-236 (**C-E**) at a multiplicity of infection (MOI) of 0.0002 and 0.02 respectively. Cells were treated or not with Iµg/ml of trypsin TPCK (dotted lines). RFe cells (dark green lines) were infected with SARS-CoV-2 (**B-D**) or BANAL-236 (**C-E**) at a MOI of 0.2 and 0.5 respectively and treated or not with Iµg/ml of tņ psin TPCK (dotted lines). (**B-C**) The relative amounts of cell-associated viral RNA were measured by RT-qPCR analysis at different times post-infection and were expressed as genome equivalents (GE) per µg of total cellular RNAs. Data arc the means ± SD of three independent experiments. (**D-E**) The percentages of cells expressing the viral N protein were determined by flow cytomctņ analysis. Data are means ± SD of three independent experiments.

Next, we examined the replication kinetics of BANAL-236 and SARS-CoV-2 (Wuhan strain) in RFe cells. Caco-2 cells, which are susceptible to both viruses [1,5] served as positive controls. Based on pilot experiments, Caco-2 cells were infected with 100 more infectious particles of BANAL-236 than SARS-CoV-2 to achieve comparable viral RNA yields. RT-qPCR analysis revealed that SARS-CoV-2 RNA levels in Caco-2 cells increased until 48 hpi, reaching about 10^7^ genome copies per μg of total RNA and remained stable through 72 hpi (Fig. 1B). The replication kinetic of BANAL-236 was similar to that of SARS-CoV-2 in Caco-2 cells (Fig. 1C). Flow cytometry analyses were used to further evaluate viral replication. Around 50% of Caco-2 cells were positive for the SARS-CoV-2 N protein at 48 hpi (Fig. 1D), while around 30% of Caco-2 cells were expressing the N protein of BANAL-236 at 72 hpi (Fig. 1E). Thus, BANAL-236 replicated less efficiently than SARS-CoV-2 in Caco-2 cells, despite the higher inoculum, a finding that contrasts with a previous study reporting comparable replication of BANAL-236 and SARS-CoV-2 (WK-521 strain) in these cells [5].

In RFe cells, infection with SARS-CoV-2 or BANAL-236 at a MOI of 0.2 and 0.5, respectively, did not result in increased viral RNA levels over time, suggesting an absence of replication (Fig. 1B-C). In agreement with these data, no N positive cells were detected by flow cytometric analysis (Fig. 1D-E). The lack of viral replication may be due to the absence of endogenous ACE2 and/or a low TMPRSS2 (Fig. 1A). To activate the S protein and allow viral fusion independently of TMPRSS2 and other proteases, viral input was treated with the serine-protease trypsin [29]. Pre-activation of SARS-CoV-2 or BANAL-236 with trypsin did not enhance viral RNA yield or the percentage of RFe cells expressing the N protein (Fig. 1B-E), suggesting that S protein cleavage is not the limiting factor for infection in RFe cells. The absence of viral RNA and protein production in RFe cells may therefore result from a lack of pro-viral factors, such as ACE2 (Fig. 1A) and/or the presence of potent antiviral mechanisms.

### Efficient entry of BANAL-236 and BANAL-52 in RFe cells requires both hACE2 and hTMPRSS2

To determine whether the absence of viral replication in RFe cells could be rescued by expressing known sarbecovirus entry factors, we generated RFe cells stably expressing human ACE2 (hA) and hTMPRSS2 (hT), or both factors (hAT), as well as 293T cells stably expressing the same factors for comparison. Flow cytometry analysis revealed that 20 to 40% of 293T cells were expressing hACE2 at their surface (Fig. 2A) while 40 to 80% were expressing hTMPRSS2 at their surface (Fig. 2B). The surface expression level of both factors was below the detection limit in wild-type cells (Fig. 2A-B). RT-qPCR analysis of the levels of exogenous hACE2 and hTMPRSS2 mRNAs in 293T cells were consistent with the flow cytometric analysis (Fig. S1A-B). We also generated 293T cells expressing *Rhinolophus* ACE2 (rA), rTMPRSS2 (rT), or both (rAT). Since BANAL-236 and BANAL-52 were isolated from *R. marshalli* and *R. malayanus*, respectively [1] and the *R. marshalli* genome is unavailable, rACE2 was synthetized from *R. malayanus* sequence. Given the endogenous TMPRSS2 expression in RFe cells (Fig. 1A), rTMPRSS2 was cloned from *R. ferrumequinum* cDNA to avoid having cells expressing TMPRSS2 from 2 species. Western blot analysis performed in 293T cells suggested that the antibodies against hACE2 recognized less efficiently rACE2 than hACE2 and that antibodies against hTMPRSS2 [30] did not detect its *Rhinolophus* ortholog (Fig. S1C).

**Figure 2.**
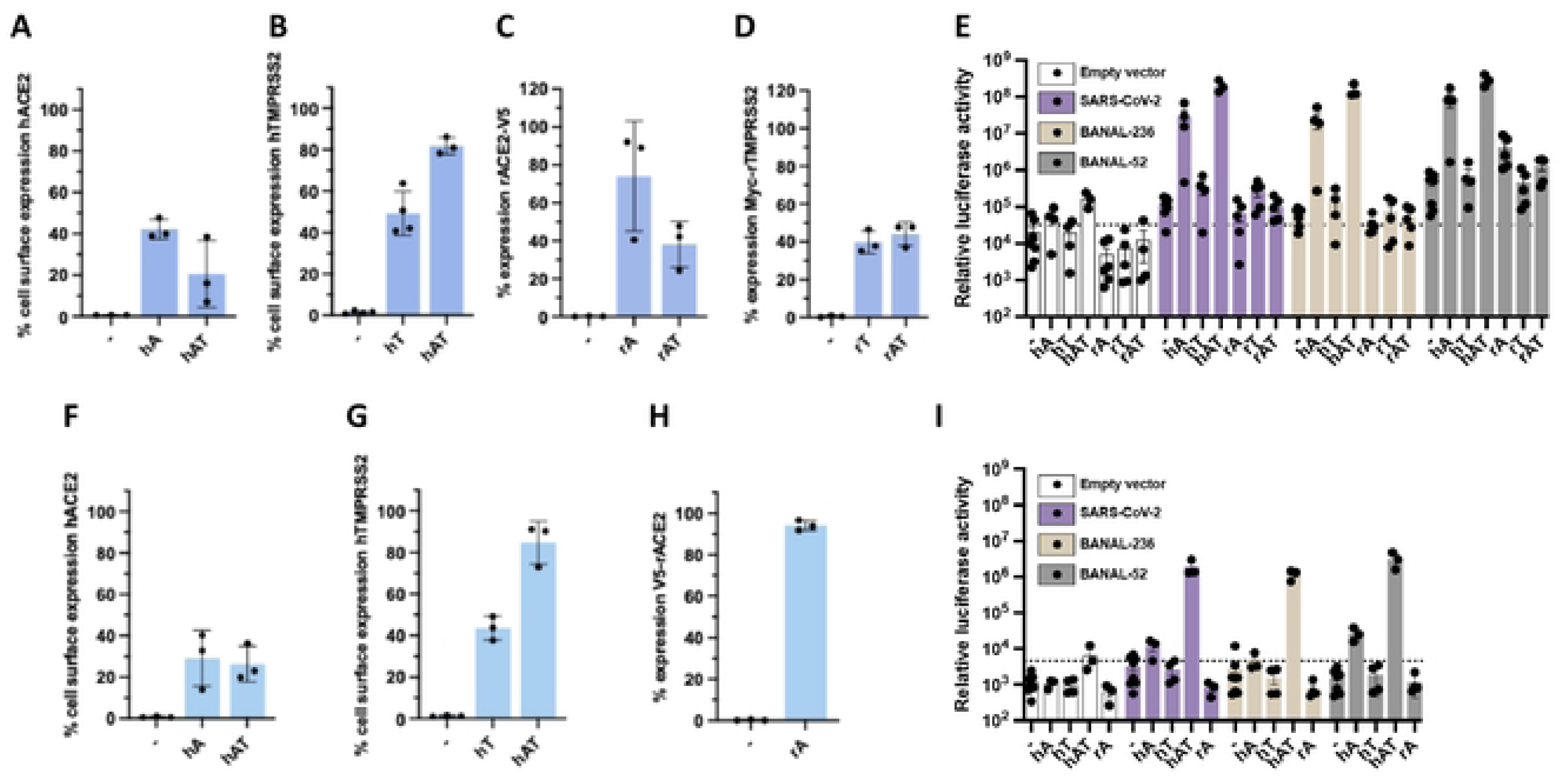
Efficient entry of BANAL-236 and BANAL-52 in RFe cells requires both hACE2 and hTMPRSS2. (**A-B**) Cell surface expression of hACE2 (**A**) and hTMPRSS2 (**B**) were determined by flow cytometry analysis prior to the entry assays shown in (**E**) in wt 293T cells (−) or expressing hACE2 (hA). hTMPRSS2 (hT) or both entry factors (hAT). Data arc the means ± SD of three independent experiments. (**C-D**). Cell surface expression of rACE2 (**C**) or rTMPRSS2 (**D**) were determined by flow cytometry analysis prior to the entry assays shown in (**E**) in wt 293T cells (−) or expressing rACE2-V5 (rA). C-myc-rTMPRSS2 (rT) or both entry factors (hAT). Data are the means ± SD of three independent experiments. (**E**) Entry assays were performed in wt 293T cells (−) or stably or transiently expressing hACE2 (hA). hTMPRSS2 (hT). rACE2-V5 (rA). C-myc-rTMPRSS2 (rT). or both entry factors (hAT or rAT). Cells were transduced with the same HIV-1 p24 quantity of pscudo-lentiviruses bearing the S proteins of SARS-CoV-2 (purple). BANAL-236 (beige) or BANAL-52 (grey). Results are expressed in relative luminescence units (RLU). Tire dashed line indicates the average RLU obtained with cells transduced with empty vectors. Data are means ± SEM of three independent experiments. (**F-H**) Cell surface expression of hACE2 (**F**), hTMPRSS2 (**G**) or rACE2 (**H**) were determined by flow cytometry analysis prior to the entry assays shown in (**I**) in wt RFe cells (−) or expressing hACE2 (hA), hTMPRSS2* (hT), both entry factors (hAT) or rACE2-V5. Data arc the means ± SD. (I) Entry assays were performed in wt RFe cells (−) or stably expressing hACE2 (hA). hTMPRSS2 (hT). both human entry factors (hAT). or rACE2 (rA). Cells were transduced with the same HIV-1 p24 quantity of pscudo-lcntiviπiscs bearing the S proteins of SARS-CoV-2 (purple). BANAL-236 (beige) or BANAL-52 (grey). Results are expressed in relative luminescence units (RLU). The dashed line indicates the average RLU obtained with cells transduced with empty vectors. Data arc the means ± SEM of three independent experiments.

Therefore, plasmids expressing tagged versions of the *Rhinolophus* entry factors (rACE2-V5 and c-myc-rTMPRSS2) were produced. Flow cytometry showed that ∼70% of 293T-rA cells expressed rACE2 and ∼40% of 293T-rAT expressed rACE2 (Fig. 2C). Around 40% of 293T-rT and 293T-rAT expressed rTMPRSS2 (Fig. 2D). RT-qPCR analysis of the abundance of rACE2 and rTMPRSS2 mRNAs in 293T cells were consistent with the flow cytometric analysis (Fig. S1D-E). ACE2 expression in 293T cells was reduced in the presence of TMPRSS2, as compared to when it was expressed on its own (Fig. 2A, 2C, S1A and S1D), likely due to the ability of the protease to cleave ACE2 [31]. Lentiviruses expressing the S proteins of SARS-CoV-2 (Wuhan strain), BANAL-236 or BANAL-52 were generated, as well as lentiviruses with no S, which served as negative controls. All three S proteins facilitated entry of pseudovirions into 293T-hACE2 cells (Fig. 2E), but not into wild-type 293T. Expression of hTMPRSS2 alone was insufficient for efficient viral entry, although co-expression with hACE2 slightly enhanced S-mediated entry (Fig. 2E). Expression of *Rhinolophus* entry factors, alone or in combination, did not support SARS-CoV-2 or BANAL-236 S-mediated entry in 293T cells (Fig. 2E), suggesting that the S protein of BANAL-236 interacts more efficiently with hACE2 than with *R. malayanus* ACE2. Expression of rACE2 alone allowed entry of lentiviruses expressing the S proteins of BANAL-52 in 293T (Fig. 2E), although not as efficiently as hACE2. This result was unexpected given that BANAL-52 was originally detected in *R*. *malayanus*. When rTMPRSS2 was expressed together with rACE2 in 293T cells, S-mediated entry of BANAL-52 was less efficient than in 293T cells expressing rACE2 alone (Fig. 2E). This is likely due to the lower expression of rACE2 in 293T-rAT cells compared to 293T-rA cells (Fig. 2C and Fig. S1D).

Flow cytometric analysis revealed that approximately 30% of RFe-hA and RFe-hAT cells expressed hACE2 at their surface (Fig. 2F), despite RFe-hAT cells expressing slightly higher levels of hACE2 mRNA than RFe-hA cells (Fig. S1F). Around 40% of RFe-hT and 90% of RFe-hAT expressed hTMPRSS2 at their surface (Fig. 2G), despite exhibiting similar abundance of hTMPRSS2 mRNA (Fig. S1G), suggesting a more efficient plasma membrane addressing of hTMPRSS2 in RFe-hAT than in RFe-hT. Finaly, about 90% of RFe-rACE2 cells expressed rACE2-V5 (Fig. 2H and S1G). Unfortunately, repeated attempts to produce RFe cells stably expressing rTMPRSS2, alone or in combination rACE2, were unsuccessful. Expression of hACE2 or hTMPRSS2 alone in RFe cells did not support efficient viral entry (Fig. 2I). Similarly, no viral entry was observed in RFe-rA cells (Fig. 2I), suggesting that expression of rACE2 alone is insufficient to mediate S-dependent entry in RFe cells, as observed in 293T cells (Fig. 2E). Only co-expression of both human entry factors enabled efficient viral entry in RFe cells (Fig. 2I). These results suggest that the undetectable levels of ACE2 and low TMPRSS2 levels in RFe cells (Fig. 1A) contribute to the absence of viral replication.

Given that RFe-hAT cells supported efficient entry of lentiviruses pseudotyped with the BANAL-236 S protein (Fig. 2I), we selected them for further investigation.

### BANAL-236 and SARS-CoV-2 replicate in RFe-ATC cells

We assessed the ability of BANAL-236, and, for comparison, SARS-CoV-2, to replicate in RFe-AT cells. Entry assays showed that these cells supported entry of lentiviruses bearing the S proteins of both viruses (Fig. 2I). However, despite maintaining RFe cells under antibiotic selection, the expression of hACE2 and hTMPRSS2 was unstable. Therefore, the cell surface expression of hACE2 and hTMPRSS2 was evaluated during each experiment by flow cytometry (Fig. 3A). Around 20% of RFe-AT cells were positive for hACE2 and 50% for hTMPRSS2 (Fig. 3A). RFe-AT cells were infected with SARS-CoV-2 or BANAL-236 at a MOI of 0.2. RT-qPCR analyzes revealed that viral RNA levels did not increase over time (Fig. 3B-C), suggesting an absence of viral replication. Although not significant, a modest increase of BANAL-236 and SARS-CoV-2 RNA was observed at late time post-infection in RFe-AT cells treated with TPCK compared to untreated cells (Fig. 3B-C), suggesting that trypsin activation of the S protein may enable a small fraction of viruses to enter cells. Consistent with the RT-qPCR results (Fig. 3B-C), flow cytometry detected no N protein expression in RFe-AT cells, even in the presence of trypsin (Fig. 3D-E). This suggests that any viruses that entered the cells were blocked at an early stage of the replication cycle. Collectively, these data demonstrate that, although hACE2 and hTMPRSS2 expression permits S-mediated entry in RFe-AT cells (Fig. 2I), it is insufficient to support viral replication.

**Figure 3.**
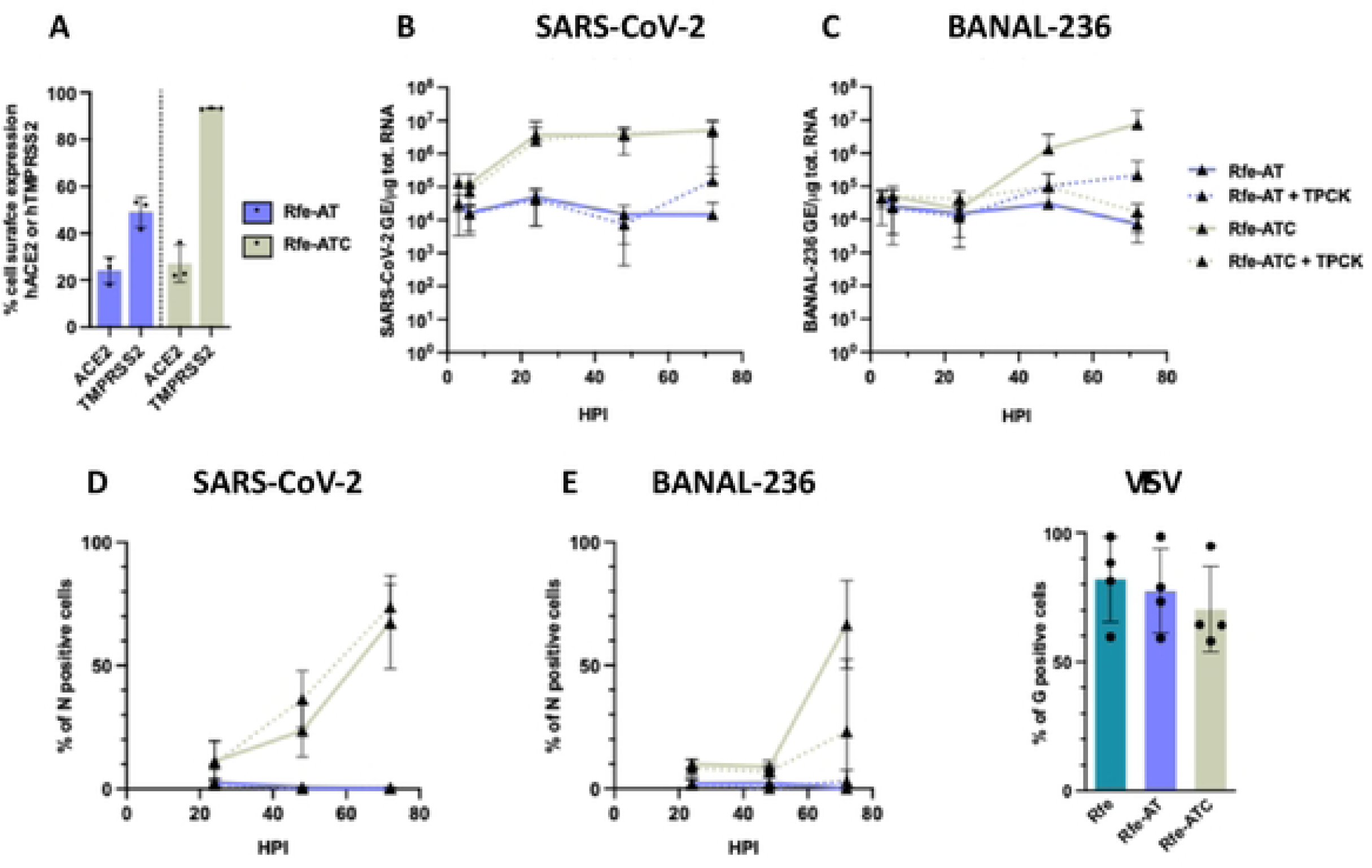
RFe-ATC cells, but not RFe cells, are permissive to SARS-CoV-2 and BANAL-236. (**A**) Expression of hACE2 at the cell surface of RFc-hACE2-hTMPRSS2 cells (RFc-AT. in blue) or clonal cells (RFe-ATC. in light green) were determined by flow cytometry analysis. Data are means ± SD of three independent experiments. RFc-AT (blue lines) and RFe-ATC (green lines) cells were infected with SARS-CoV-2 (**B-D**) or BANAL-236 (**C-E**) at a multiplicity of infection (MOI) of 0.2 and 0.5. treated or not with Iµg/ml of trypsin TPCK (dotted lines). Cells w ere collected at the indicated hours post-infection (HPI). (**B-C**) The relative amounts of cell-associated viral RNA were measured by RT-qPCR analysis and were expressed as genome equivalents (GE) per µg of total cellular RNAs. Data arc the means ± SD of three independent experiments. (**D-E**) The percentages of cells expressing the viral N protein were determined by flow cytometry analysis. Data arc means ± SD of three independent experiments. (**F**) RFe. RFc-AT and RFe-ATC cells were infected w ith VSV at a MOI of 0.05 for 16 hours. The percentages of cells expressing the viral G protein were determined by flow cytometry analysis. Data arc means ± SD of three independent experiments.

Due to the unstable expression of hACE2 and hTMPRSS2 in RFe-AT cells, we produced over 30 clonal cell lines. Among these, a single clone (RFe-ATC) exhibited a cytopathic effect following SARS-CoV-2 infection. Flow cytometry analysis revealed that around 25% of RFe-ATC cells were positive for hACE2 and 90% for hTMPRSS2 (Fig. 3A). RFe-ATC cells were infected with SARS-CoV-2 or BANAL-236 at a MOI of 0.2 and 0.5, respectively. RT-qPCR analyzes showed that SARS-CoV-2 RNA levels increased by approximately 1-log during the first 20 hours of infection, followed by a plateau (Fig. 3B). In contrast, BANAL-236 RNA levels rose steadily between 20– and 72-hours post-infection in RFe-ATC cells (Fig. 3C). Flow cytometry analysis showed that a small percentage of RFe-ATC cells were positive for SARS-CoV-2 N protein at 24 hpi, increasing to 60% at 72 hpi (Fig. 3D). Consistently with the viral RNA yield data (Fig. 3C), BANAL-236 replication in RFe-ATC cells was delayed compared to SARS-CoV-2 (Fig. 3E). Taken together, these results demonstrate that both SARS-CoV-2 and BANAL-236 are capable of replicating in RFe-ATC cells.

To determine whether RFe parental and RFe-AT were generally resistant to viral infection, we infected them with Vesicular Stomatitis virus (VSV), a negative-strand RNA virus from the *Rhabdoviridae* family that is known for its broad cell tropism [32]. Flow cytometry analysis using an antibody against the VSV G protein revealed that VSV replicated similarly in RFe, RFe-AT and RFe-ATC cells (Fig. 3F), suggesting that the inability of SARS-CoV-2 and BANAL-236 to replicate in RFe and RFe-AT cells is not due to a pre-existing antiviral state.

The high surface expression of hTMPRSS2 in RFe-ATC cells (Fig. 3A) may underlie their susceptibility to SARS-CoV-2 and BANAL-236. To test this hypothesis, we used siRNAs to knock-down hTMPRSS2 expression, selecting a concentration that reduced the proportion of hTMPRSS2-positive RFe-ATC cells (Fig. 4A), to levels comparable to those in RFe-AT cells (Fig. 3A). Under these conditions, BANAL-236 and BANAL-52 S-mediated entry was modestly, but reproducibly, less efficient in RFe-ATC cells expressing reduced levels of hTMPRSS2, as compared to cells transfected with control siRNAs (Fig. 4B). Viral replication in the presence of reduced levels of hTMPRSS2 was then assessed by flow cytometry by measuring the number of N protein-positive cells at 72 hpi. Viral replication was completely abolished in these cells, as compared to cells treated with control siRNAs (Fig. 4C), suggesting a strong dependence on hTMPRSS2. This pronounced effect likely reflects the cumulative impact of reduced hTMPRSS2 expression over multiple rounds of replication, in contrast to the modest effect observed during a single-entry event. Thus, the susceptibility of the RFe-ATC cells to viral infection may be attributed to their high level of hTMPRSS2 expression, as compared to RFe-AT cells.

**Figure 4.**
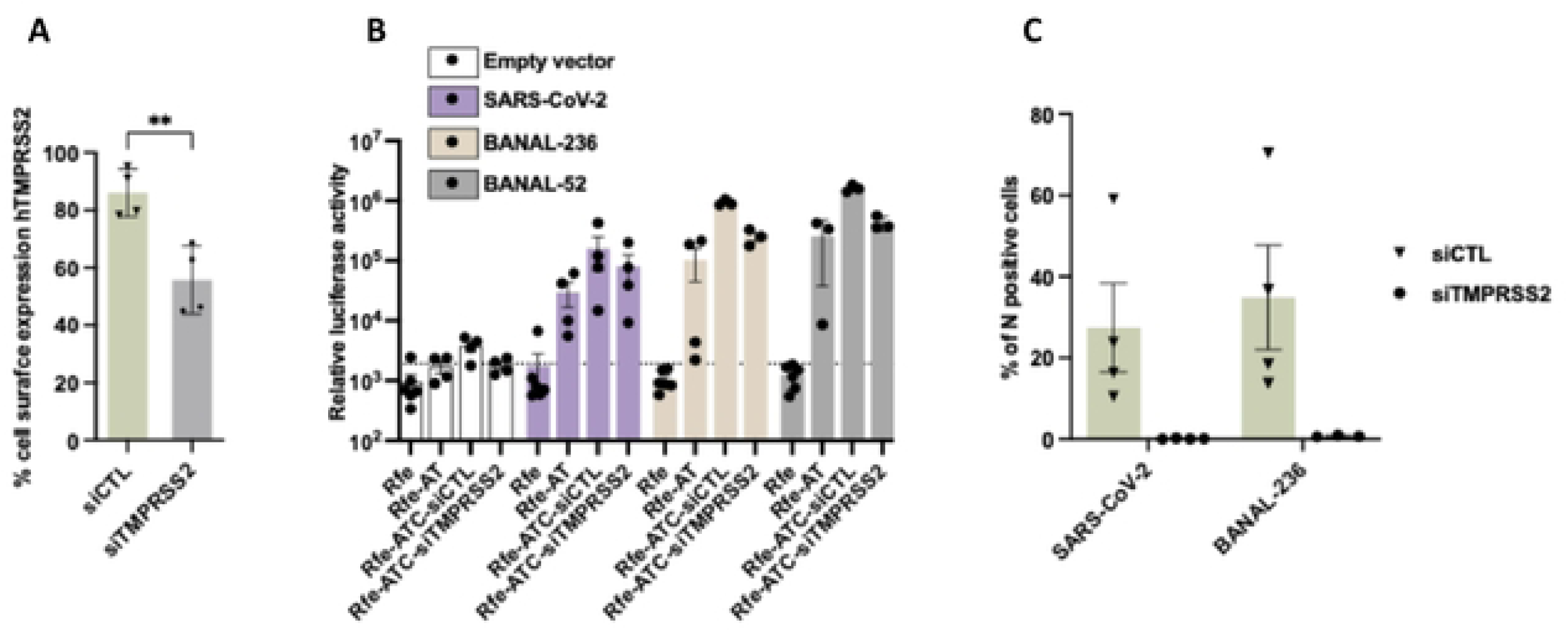
BANAL-236 replication in RFe-ATC cells is dependent on high cell surface TMPRSS2 expression. (**A**) RFe-ATC cells were transfected with siRNAs targeting TMPRSS2 or control siRNAs (siCTL) for 3 days. The percentages of cells expressing hTMPRSS2 at their surface were determined by flow cytometry analysis. Data are the means ± SD of four independent experiments. T-tests two-tailed were performed p<().OI. (**B**) Entry assays were performed with Icntiviruscs pseudotyped with the S proteins of SARS-CoV-2 (purple). BANAL-236 (beige) or BANAL-52 (grey). Cells were transduced with the same HIV-1 p24 quantity of pscudo-lcntiviπiscs. Results arc expressed in relative luminescence units (RLU). The dashed line indicates the average RLU obtained with cells transduced with empty vectors. (**C**) Cells were infected with SARS-CoV-2 and BANAL-236 at a multiplicity of infection (MOI) of 0.02 and 0.2. respectively. Seventy-two hours later, cells were collected and the percentages of cells expressing the viral N protein were determined by flow cytometry analysis. Data arc means ± SD of at least three independent experiments.

### SARS-CoV-2 and BANAL-236 complete their replication cycles in RFe-ATC cells

To further characterize SARS-CoV-2 and BANAL-236 replication in RFe-ATC cells, we performed transmission electron microscopy on cells infected for 72 h, with Caco-2 cells included for comparison and non-infected cells as negative controls (Fig. S2). In infected Caco-2 cells, we observed numerous viral replication factories composed of double-membrane vesicles (DMVs) (Fig. 5A-B), as previously described in SARS-CoV-2 infected Vero cells [33], although the DMVs in these cells displayed slight structural differences consistent with variations reported across different cell lines [34]. These DMVs, which likely derive from ER membranes and autophagic processes, serve as sites for viral RNA synthesis [34,35]. Fully assembled virus particles were detected within large intracellular vesicles (Fig. 5C), some of which fused with the plasma membrane (Fig. 5D), suggesting virion release via exocytosis. Many virions remained attached to the surface of infected cells (Fig. 5D), possibly due to high hACE2 expression at the plasma membrane. RFe-ATC cells infected with SARS-CoV-2 exhibited similar ultrastructural changes (Fig. 5E-H), including numerous DMVs. Large vacuoles containing virions (Fig. 5G) and surface-bound virions (Fig. 5H) were also present in RFe-ATC cells. In contrast, Caco-2 and RFe-ATC cells infected with BANAL-236 contained very few DMVs (Fig. 5I-P), with only occasional DMV-like structures observed (Fig. 5N, white arrows). In addition, large vacuoles filled with dozens of BANAL-236 virions were observed in both cell lines (Fig. 5J and 5O), some of which were connected to the plasma membrane (Fig. 5P). The presence of virions at the surface of RFe-ATC cells (Fig. 5N and 5P) suggested that BANAL-236 completed its replication cycle in these bat cell line.

**Figure 5.**
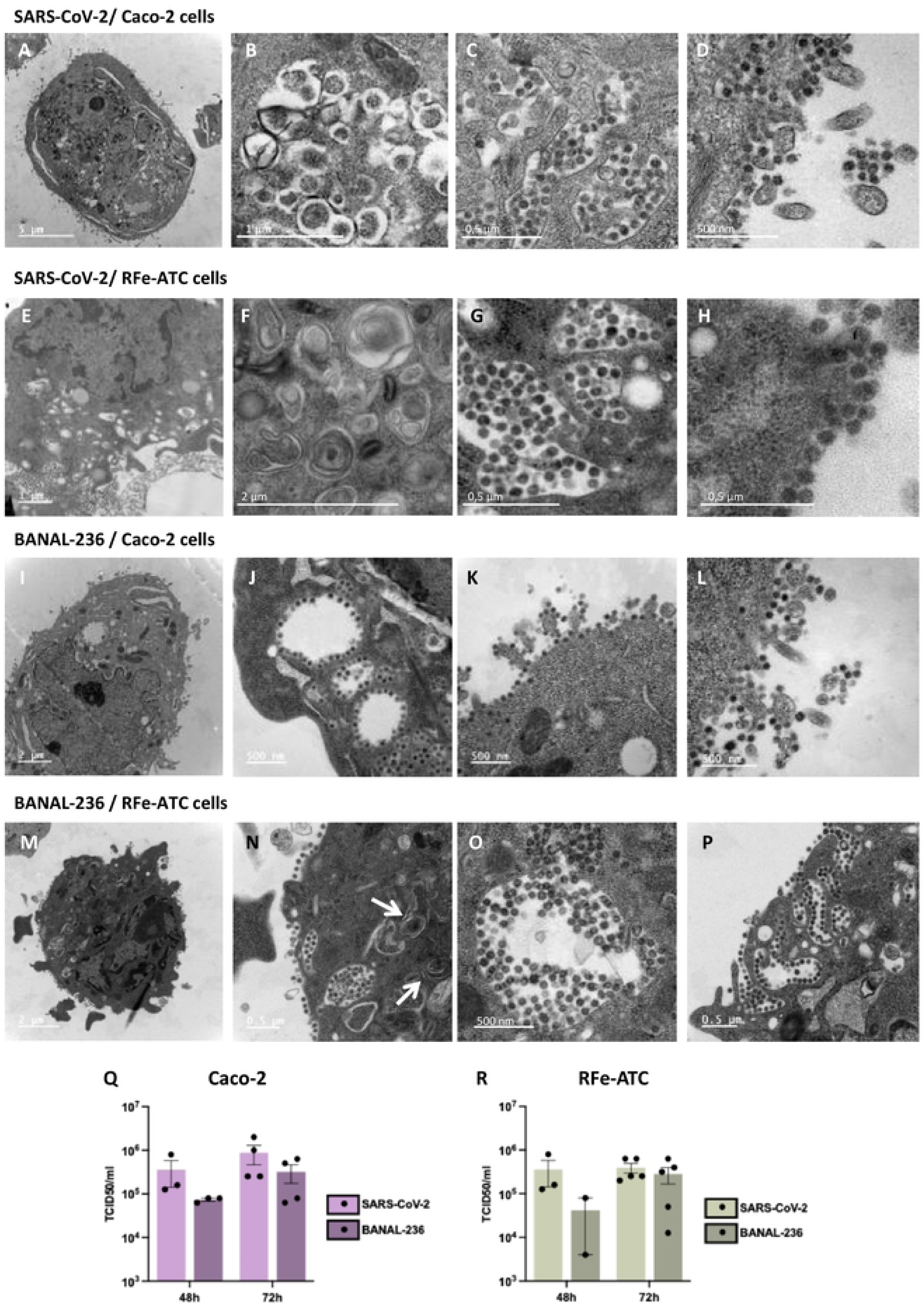
SARS-CoV-2 and BANAL-236 complete their replication cycle in RFe-ATC cells. (**A-P**) Caco-2 and RFe-ATC cells were infected for 72 h. Caco-2 cells were infected with SARS-CoV-2 (**A-D**) or BANAL-236 (**I-L**) at an MOI 0.001 and 0.01 respectively. RFe-ATC cells were infected with SARS-CoV-2 (**E-H**) and BANAL-236 (**M-P**) at MOI 0.02 and 0.2 respectively. Cells were then subjected to transmission electron microscopy analysis. DMV-likc structures arc pointed by white arrows. (**Q-R**) At 4X and 72 h post-infection supernatants from Caco-2 (**Q**) and RFe-ATC (**R**) cells were collected. Titration of clarified supernatants by TCIDjo assays was performed on VERO-E6 cells infected for 5 days. Data arc the means ± SEM of at least two independent experiments.

To assess the infectivity of virions observed at the cell surface, supernatants from infected RFe-ATC and Caco-2 cells were titrated on Vero-E6 cells (Fig. 5Q-5R). Supernatants from Caco-2 cells contained approximately 5.10^5^ TCID_50_/mL of infectious BANAL-236 or SARS-CoV-2 particles (Fig. 5Q). Similarly, about 10^5^ TCID_50_/mL were retrieved from the supernatant of infected RFe-ATC cells (Fig. 5R). These results confirmed that both viruses complete their replication cycles in RFe-ATC cells.

### BANAL-236 antagonizes the IFN response in *Rhinolophus* and human cells

Given the critical role of viral IFN antagonism in overcoming species barrier, we evaluated ISG induction in RFe cells and their derivatives following infection with BANAL-236 or SARS-CoV-2. We compared mRNA abundance of *OAS1* and *ISG20*, two ISGs that are conserved across vertebrate species [36], in infected cells. No upregulation of *OAS1* or *ISG20* expression was observed in RFe nor RFe-AT cells at 72 hpi (Fig. 6A-B), which was consistent with the absence of viral replication in these cells (Fig. 1 and 3). Despite robust replication of BANAL-236 and SARS-CoV-2 in Caco-2 and RFe-ATC cells (Fig. 1 and 3), no induction of *OAS1* or *ISG20* expression was observed either at 72 hpi (Fig. 6A-B). Similar results were obtained when the mRNA abundance of these two ISGs were evaluated in cells infected for 4, 24, and 48 hours (Fig. S3). These results suggest that both viruses efficiently counteract the IFN response in these cell lines. To ensure that RFe cells were immunocompetent, we assessed ISG induction upon stimulation with poly I:C, a synthetic dsRNA analog. The mRNA levels of *OAS1* and *ISG20* increased upon stimulation in all four cell lines (Fig. 6A-B), demonstrating intact IFN-induction and –signaling pathways.

**Figure 6.**
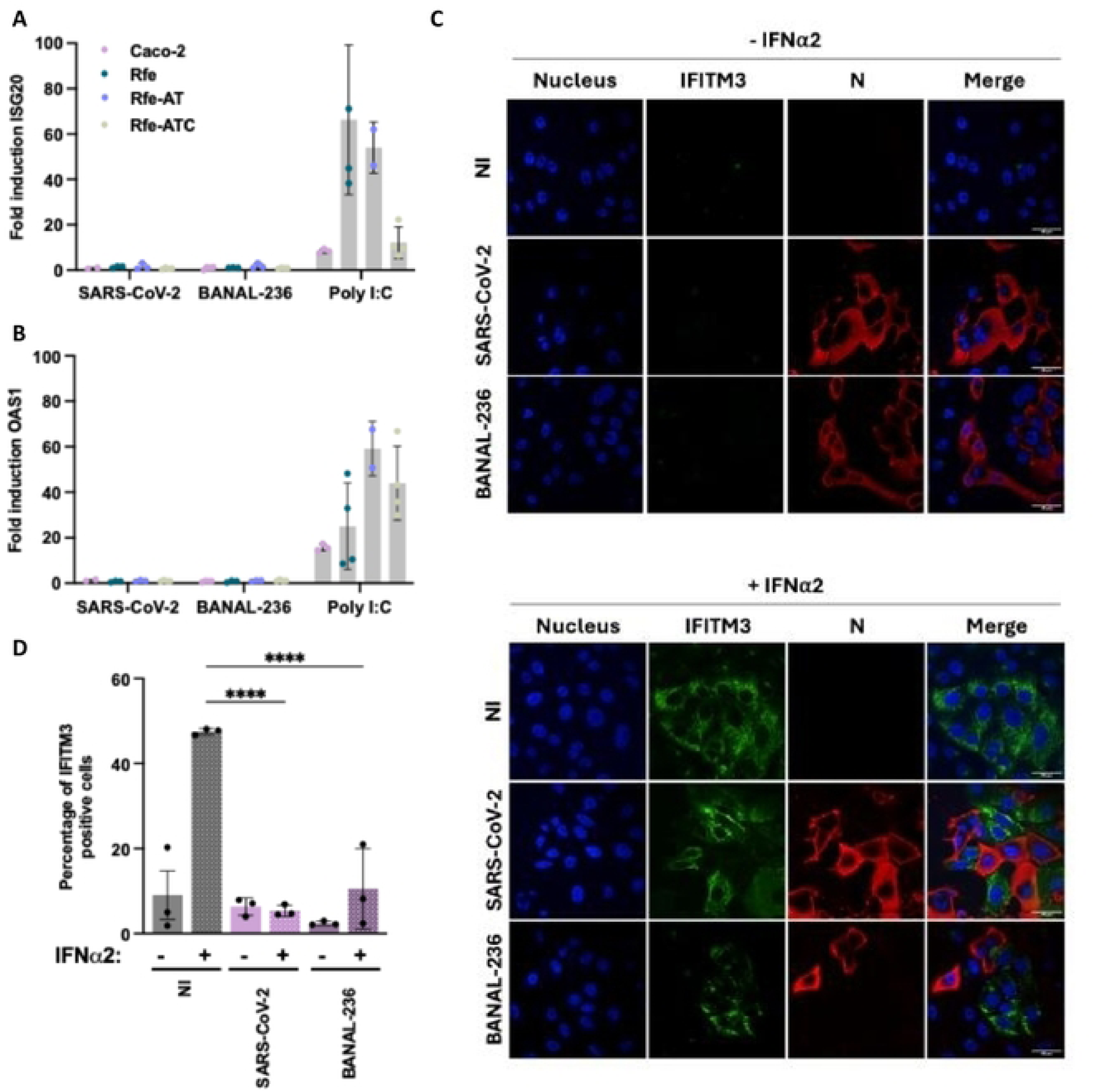
SARS-CoV-2 and BANAL-236 replication failed to induce the expression of Interferon-stimulated genes in Caco-2 and RFe-ATC cells. (**A-B**) Caco-2 (pink). RFc (dark green). RFe-AT (blue) and RFe-ATC (grey) cells were infected with SARS-CoV-2 or BANAL-236 at a multiplicity-of infection (MOI) of 0.2 and 0.5. respectively, for 72 hours. Alternatively, cells were treated with 100-500 ng of poly I:C for 16 hours. The relative amounts of ISG20 (**A**) and OASI (**B**) mRNAs were determined by RT-qPCR analysis. Results were first normalized to GAPDH mRNA and then to mRNA levels of control cells (mock-treated or non-treated cells), which were set at I. Data are means ± SD of at least two independent experiments. (C) Caco-2 cells were infected for 72 hours with SARS-CoV-2 or BANAL-236 at a MOI of 0.0002 and 0.02. respectively, or left non-infected (Nl). They were stimulated with IFNα2 at I OOOIU/ml (lower panel) or mock-treated (upper panel) 16 hours before fixation. They were stained with antibodies recognizing IFITM3 (green) and the viral nuclcocapsid N (red). Nucleus were counterstained using Hoechst dye (blue). Images are representative of three independent experiments. Scale bars. 40 µm. (D) Percentages of IFITM3 – positive cells among cells expressing the viral N proteins were estimated by analyzing at least 40 cells per condition. For uninfected (Nl) cells, at least 60 cells were randomly picked. Data are mcan ± SD. One-way ANOVA tests with Dunnell’s correction were performed. ****P< 0.0001.

To further investigate the ability of the 2 viruses to antagonize IFN signaling in individual cells, we assessed IFITM3 expression, another ISG that is conserved across vertebrate species [36], using fluorescent microscopic assays in Caco-2 cells (due to the lack of suitable antibodies for *Rhinolophus* ISGs). IFITM3 was detected in about 10% of unstimulated Caco-2 cells (Fig. 6C-D) but in approximately 45% of cells treated with IFN⍺2 (Fig. 6C-D). In cells infected for 72 hours, IFITM3 was detected in fewer than 5% of N-positive cells (Fig. 6C-D). When infected cells were treated with IFN⍺2 16 hours before fixation (Fig. 6C-D), fewer than 10% of cells were positive for both N and IFITM3, confirming that SARS-CoV-2 or BANAL-236 replication limits ISG expression, as observed in the RT-qPCR analysis (Fig. 6A-B).

To identify BANAL-236 proteins capable of antagonizing the IFN response in human cells, the open reading frames (ORFs) of all 30 viral proteins were cloned downstream of a N-terminal 3X-FLAG tag and expressed in 293T cells, which are easy-to-transfect cells. Western blot analyses confirmed the expression of 27 BANAL-236 proteins at their expected sizes (Fig. 7A), including NSP3, the largest coronavirus protein (∼220 kDa). A second ∼80 kDa NSP3 band was detected, as previously reported for SARS-CoV-2 Nps3 [19]. NSP3 is composed of around 15 domains, including the papain-like protease [37], and this extra band could represent an autoproteolytic product. FLAG-ORF8 was poorly expressed, and FLAG-S was unproperly processed. We thus generated ORF8 and S sequences with a C-terminal STREP-tag. ORF8-STREP was detected at the correct size (Fig. 7A). Both the uncleaved precursor and the dissociated S2 fragment of S-STREP were detected, as expected [38]. Due to its small size (1.5 kDa), NSP11 was undetectable and was thus excluded from the analysis.

**Figure 7.**
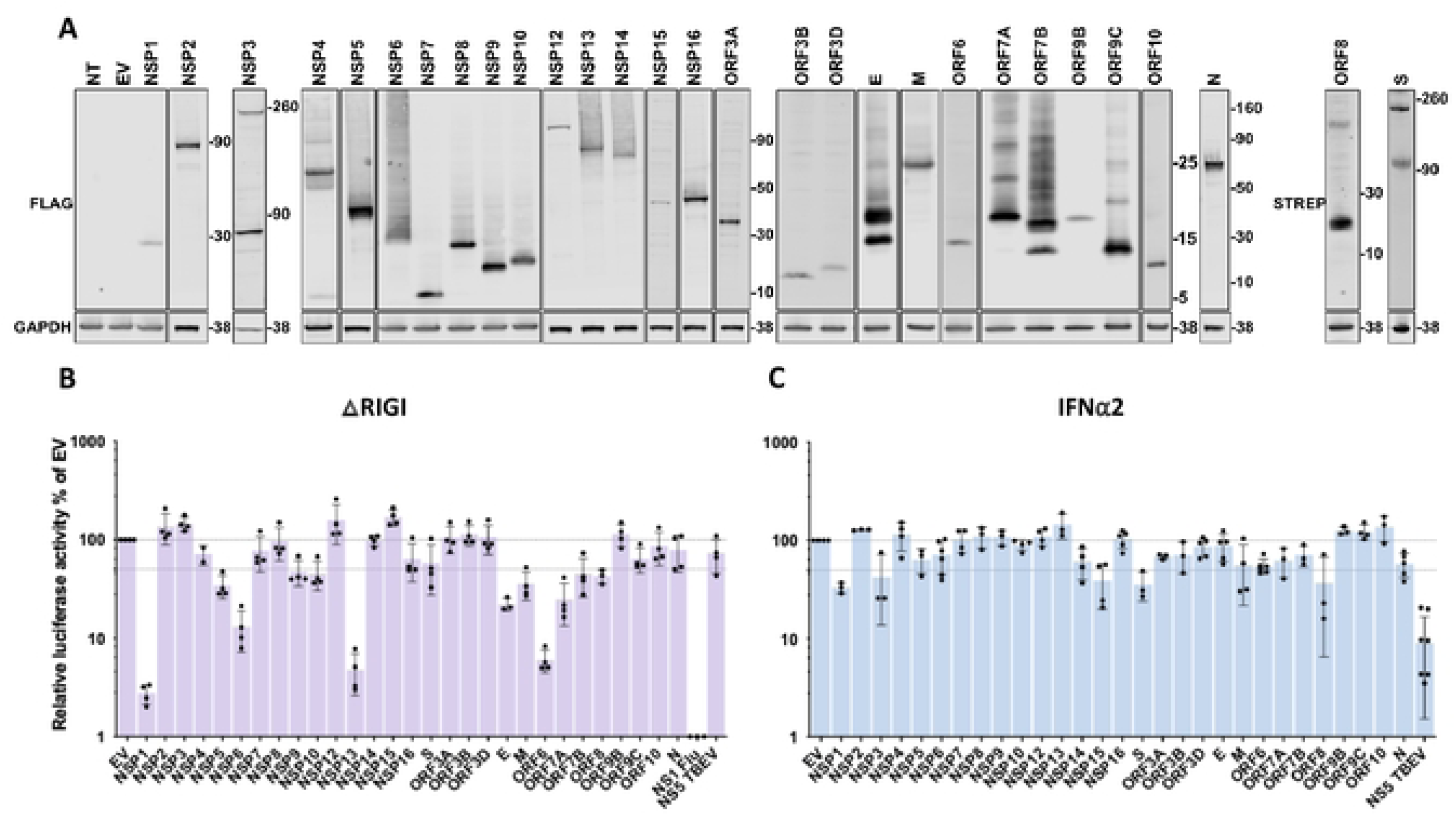
BANAL-236 antagonizes the 1FN response in human cells. (**A**) Viral protein expression in 293T cells. 293T cells were mock – transfected (NT), transfected with empty plasmids (EV) or with plasmids encoding FLAG or STREP – tagged BANAL-236 viral proteins. Cells were harvested 24 h post – transfection and protein expression was assessed by Western blotting with anti – FLAG. anti-STREP and anti – GAPDH antibodies. Data arc representative of three biological replicates. (**B**) 293T cells were co – transfected for 24h with the Firefly luciferase reporter plasmid p – 1SRE – luc. TK Rcnilla luciferase control plasmid phRluc – TK. ΔRIGI plasmid and plasmids encoding individual viral proteins of BANAL-236. Empty vectors (EV) and plasmids encoding the NS I of Flu protein were used as negative and positive controls, respectively. The data were analyzed by first noπnalizing the Firefly luciferase activity to the Rcnilla luciferase (Rluc) activity and then to EV samples, which were set at 100%. Data arc means ± SD of at least three biological replicates. (**C**) 293T cells were co – transfected with the Firefly luciferase reporter plasmid p – 1SRE – luc. TK Rcnilla luciferase control plasmid phRluc – TK, and plasmids encoding individual viral proteins of BANAL-236. Empty vectors (EV) and plasmids encoding the NS5 of TBEV protein were used as negative and positive controls, respectively. Cells were stimulated 7 h post – transfection with IFNα2 at 200 IU/ml and assayed for luciferase activity 16 h later. The data were analyzed by first noπnalizing the Firefly luciferase activity to the Renilla luciferase (Rluc) activity and then to EV samples, which were set at 100%. Data arc means ± SD of at least three biological replicates.

The effect of BANAL-236 proteins on the activation of the IFN-stimulated response elements (ISREs) was investigated in 293T cells overexpressing a constitutively active form of RIG-I (ΔRIG-I), a potent inducer of IFN production [39]. ISREs can be activated directly by transcription factors of the IFN induction pathway, like IRF3, in an IFN-independent manner [40]. Cells transfected with an empty plasmid (EV) served as references. Cells expressing a FLAG-tagged version of the NS1 of influenza A virus (IAV), a known IFN antagonist [41], served as positive controls. Its expression indeed reduced ISRE activation by over 100-fold (Fig. 7B). TBEV NS5, which inhibits the JAK-STAT pathway [42], had no effect on ISRE activation (Fig. 7B), confirming that ISRE was activated in an IFN-independent manner in our experimental settings. Four BANAL-236 proteins diminished the activity of the ISRE by at least10-fold: NSP1, NSP6, NSP13, and ORF6 (Fig. 7B), mirroring the antagonistic effects of their SARS-CoV-2 counterparts detected in similar experiments [16,17,20], despite sequence differences (Table S1). To determine whether BANAL-236 proteins were capable of inhibiting IFN-I signaling, we repeated the screen in 293T cells stimulated with IFN⍺2 (Fig. 7C). As expected [42], TBEV NS5 inhibited ISRE activation by around 10-fold (Fig. 7C). NSP1, NSP3, NSP15 and S modestly suppressed the JAK-STAT pathway (Fig. 7C). By contrast, SARS-CoV-2 expresses several proteins that potently block ISRE activation in IFN-I treated 293T cells, such as NSP1, NSP6, NSP13, NSP14, ORF6 and ORF7b [16,17,19,20]. Together, these findings showed that BANAL-236 is armed with mechanisms to evade the IFN response in human cells, albeit with a potential narrower repertoire of IFN-I signaling antagonists compared to SARS-CoV-2.

## Discussion

We established a fibroblast line from the lung tissue of *R. ferrumequinum* (RFe), a bat species distributed across Europe, Northern Africa, Central Asia and Eastern Asia. These cells were refractory to infection with BANAL-236 and SARS-CoV-2 replication. They did not allow entry of lentiviruses pseudotyped with the S protein of either of these two sarbecoviruses, or of BANAL-52. This resistance aligns with the absence of ACE2 expression and low levels of TMPRSS2 in these cells. RFe-AT cells, which stably expressed hACE2 and hTMPRSS2, were also resistant to viral replication, despite supporting entry of S-bearing lentiviruses. Overcoming the entry blockade in RFe-AT cells was thus insufficient to permit viral replication, suggesting that post-entry mechanisms, such as the absence of essential pro-viral factors and/or the presence of potent antiviral factors, inhibit the replication of sarbecoviruses in these cells. Only one clonal cell line derived from RFe-AT cells (RFe-ATC) was permissive to BANAL-236 and SARS-CoV-2. A high proportion of these cells expressed hTMPRSS2 at their surface. Our results suggest that this elevated hTMPRSS2 expression may underlie their permissiveness, as even a modest reduction in hTMPRSS2 levels in RFe-ATC cells abolished viral replication over 3 replication cycles. This is consistent with studies in human cells, where TMPRSS2-mediated cell surface fusion enhances SARS-CoV-2 entry and downstream events, such as viral replication, transcription, and release [43]. Our data suggest a similar role for TMPRSS2 in facilitating BANAL-236 replication in RFe-ATC cells. Additional differences between RFe-AT and RFe-ATC cells, such as variations in basal expression of IFNs or antiviral ISGs, may further explain their distinct permissiveness to viral replication.

The receptor-binding domain (RBD) of the S protein of sarbecoviruses is a critical determinant of ACE2 binding and, consequently, viral host range. Among the 17 ‘contact residues’ essential for ACE2 interaction [44–46], the RBDs of BANAL-52 and –236 differ from that of SARS-CoV-2 by only one (H498Q) or two (K493Q and H498Q) residues, respectively [1]. Given this high sequence similarity, we anticipated that these viruses would bind with high affinity to ACE2 from their natural hosts and closely related species. However, S-mediated entry of BANAL-236 and BANAL-52 was more efficient in 293T cells expressing hACE2 than in 293T cells expressing comparable levels of *R. malayanus* ACE2. This trend extended to a broader panel of *Rhinolophus* ACE2 orthologs. Entry of lentiviruses carrying the S proteins of BANAL-236 and BANAL-52 was more efficient in 293T expressing hACE2 than in 293T cells expressing ACE2 from a 7 other *Rhinolophus* species (*R. affinis*, *R. cornutus*, *R. ferrumequinum*, *R. macrotis*, *R. pearsonii*, *R. pusillus*, *R. shameli*, and *R. sinicus)* [47]. Although ACE2 expression levels were not quantified in these experiments [47], these results, together with ours, suggest the S proteins of BANAL-236 and BANAL-52 bind hACE2 with higher affinity than ACE2 from at least 8 *Rhinolophus* species, including *R*. *malayanus*, the species from which the BANAL-52 sequence was recovered [1]. This unexpected binding preference raises intriguing questions. BANAL-52 may circulate in multiple *Rhinolophus* species and its reservoir host could be a species other than *R*. *malayanus*. Supporting this possibility, lentiviruses bearing the BANAL-52 S protein entered 293T cells more efficient when expressing ACE2 from *R. macrotis*, *R. shameli* and *R. sinicus*, compared to cells expressing the other 4 tested *Rhinolophus* ACE2 [47]. One of these 3 species could thus be the natural reservoir for BANAL-52. To further clarify the host range of BANAL-related coronaviruses, future studies should systematically evaluate the binding affinity of their RBDs to ACE2 from a comprehensive collection of *Rhinolophus* species.

The inability of rACE2 to facilitate viral entry for BANAL-CoVs could be explained by the usage of alternative receptors in *Rhinolophus* cells. Entry assays using the S proteins from clade 2 sarbecoviruses, including those isolated in *R. sinicus,* showed ACE2-independent entry into *R. sinicus* intestinal primary cells [48]. Similarly, RsHuB2019, another clade 2 sarbecovirus isolated from *R. sinicus* fecal samples, entered both human and bat cells without relying on ACE2 [49]. ACE2-independent entry has also been reported for SARS-CoV-2 in several cellular models, such as myeloid cells [50,51] and human H522 lung adenocarcinoma cells [52]. Thus, more investigations are needed to study receptor usage of BANAL-CoVs in relevant bat cells.

Our findings reveal that BANAL-236 replicated less efficiently than SARS-CoV-2 in RFe-ATC cells. This attenuated replication aligns with previous reports in human primary lung cells and in hamster lungs [4]. The lower replication rate is not due to poor receptor affinity: BANAL-236’s S protein binds hamster and human ACE2 with even higher affinity than the S protein of early SARS-CoV-2 isolates [1,5]. The attenuated growth and transmission of BANAL-CoVs, as compared to SARS-CoV-2, may partially result from the absence of an FCS. This hypothesis is supported by studies showing that SARS-CoV-2-ΔFCS exhibits reduced pathogenicity in hACE2-mice and hamsters when intranasally inoculated [5,53], and lower transmissibility in ferrets, compared to wild-type SARS-CoV-2 [54]. However, since BANAL-236 replicates even less efficiently than SARS-CoV-2-ΔFCS in human iPSC-derived airway epithelial cells and hamsters [5], the lack of an FCS is not the sole determinant of its attenuated growth in human respiratory cells. Additionally, the low percentage of N-positive RFe-ATC cells observed up to 48 hpi, suggests that BANAL-236 may have acquired adaptive mutations to enhance its replication after two rounds of infection.

BANAL-236 was isolated from bat feces and could thus be an enterotropic virus in *Rhinolophus* bats [1]. Consistently, in macaques infected *via* the nasal and tracheal routes simultaneously, BANAL-236 behaved like an enteric virus [6]. Furthermore, BANAL-236 replicated more efficiently than SARS-CoV-2 in iPSC-derived human colon organoids, particularly in colonocytes [5]. Like SARS-CoV and SARS-CoV-2 in humans [55–57], BANAL-236 may infect both the respiratory and gastrointestinal tracts of *Rhinolophus* bats. It would be of interest to generate *Rhinolophus marshalli* cell lines derived from gastrointestinal tissues for further investigations. Experimental infection of available bat colonies with BANAL-236 and/or BANAL-52 could also provide further insight into the tropism of BANAL-CoVs in bat species.

Kinetic experiments performed in Caco-2 or RFe-ATC cells revealed that BANAL-236 replication failed to induce *ISG20* and *OAS1,* despite responding well to poly I:C stimulation. Consistently, BANAL-236 and BANAL-52 infection triggered only modest ISG induction in primary human bronchial and nasal epithelial cells [4]. These results suggest that both viruses suppress IFN responses across several cellular models. Additionally, BANAL-236 and BANAL-52 were resistant to IFN treatment in Vero-hACE2-hTMPRSS2 cells [4], indicating that they have evolved potent mechanisms to evade immune response in diverse mammalian cells. This is not surprising since SARS-CoV-2 efficiently counteracts IFN response through many of its proteins [16,20] and that BANAL-236 shares about 99.96 % nucleotide sequence identity with SARS-CoV-2 (Table S1). We identified four BANAL-236 proteins (NSP1, NSP6, NSP13, and ORF6) that potently antagonize the IFN induction pathway in human cells. Their SARS-CoV-2 counterparts are known to be IFN antagonists [16,17,19,20]. However, we could not assess the IFN-evasion capacity of these proteins in RFe cells due to their low transfection and transduction efficiency. While BANAL-CoVs likely never encountered human cells, they have evolved to evade IFN responses in their natural bat hosts and the components that they target are probably highly conserved across mammalian species. For example, SARS-CoV-2 NSP1 and NSP13 prevent the phosphorylation of human IRF3 and TBK1 respectively [58,59]. BANAL-236 NSP1 and NSP3 likely employ similar mechanisms to target IRF3 in *Rhinolophus* cells and, consequently, in human cells. Our findings suggest that BANAL proteins more efficiently antagonize components of the IFN induction pathway than those of the JAK/STAT pathway, possibly because the JAK/STAT pathway is more divergent between humans and bats.

In summary, our findings show that BANAL-236, the likely ancestor of SARS-CoV-2, possesses key features that facilitated zoonotic spillover: its entry can be mediated by human entry factors and it efficiently evades human IFN responses. Additionally, the RFe-ATC cells we developed provide a valuable new model for studying molecular interactions between sarbecoviruses and their natural hosts.

## Materials and Methods

### Cell lines

Human colorectal adenocarcinoma Caco-2 cells (kind gift from Nathalie Sauvonnet, Institut Pasteur, Paris), African green monkey kidney epithelial Vero-E6 cells (ATCC CRL-1586), and human embryonic kidney (HEK) 293T cells (ATCC CRL-3216), hereinafter referred to as 293T, were maintained in Dulbecco’s Modified Eagle Medium (DMEM) (Gibco) containing GlutaMAX I, sodium pyruvate (Invitrogen) supplemented with 10% heat-inactivated fetal bovine serum (FBS) (Dutscher) and 1% penicillin and streptomycin (10 000 IU/ml; Thermo Fisher Scientific). The *Rhinolophus ferrumequinum* cells (RFe) were derived from lung tissue of a bat captured in Jerez, in Southern Spain. Primary lung fibroblasts were immortalized using lentiviruses expressing the Simian Vacuolating Virus 40 large T antigen (SV40T)[28]. RFe cells were cultured in medium containing 50% DMEM with GlutaMAX I, sodium pyruvate (Invitrogen) and 50% F-12 (Ham) medium (Gibco), supplemented with 10% heat-inactivated fetal bovine serum (FBS) (Dutscher) and 1% penicillin and streptomycin (10.000 IU/ml; Thermo Fisher Scientific). All cells were maintained at 37°C in a humidified atmosphere with 5% CO_2_.

RFe and 293T cell lines stably expressing hACE2, hTMRSS2, rACE2-V5 or C-myc-rTMPRSS2 were generated by lentiviral transduction. 2.10^5^ cells were seeded in 6-well plates. Cells were transduced with 300 μL of lentivirus in 1 ml of medium containing 1x Polybrene the following day. After 3 days of incubation at 37°C, cells were either directly used for experiments or placed under antibiotic selection (6 μg/ml of puromycin and/or 6 μg/ml of blasticidin for RFe cells; 1 μg/ml of puromycin and 10 μg/ml of blasticidin for 293T cells). RFe cells stably expressing hACE2 and hTMPRSS2 (RFe-AT) were selected with 5 μg/ml blasticidin and 250 μg/ml hygromycin (Invitrogen). RFe-AT cells were used to derivate clones (RFe-ATC) using limiting dilution.

### Viruses and infections

Experiments with BANAL-236 and SARS-CoV-2 viruses were conducted in a Biosafety Level 3 (BSL-3) laboratory, following safety protocols approved by the Institut Pasteur’s risk prevention service. The SARS-CoV-2 strain BetaCoV/France/IDF0372/2020 and BANAL-236 were obtained from the National Reference Center for Respiratory Viruses hosted by Institut Pasteur. Viral stocks were produced by amplification on Vero-E6 cells, for 72 h in DMEM supplemented with 2% FBS and 1% P/S.

Cells were exposed to the virus for 3 hours in a low volume of FBS-free medium. Cells were infected with the viruses at different MOIs as indicated in each figure. For infection in the presence of TPCK-trypsin, it was added after 3 hours of infection, at a concentration of 1 μg/ml, in the medium described above or in DMEM containing 2% FBS. Experiments with Vesicular stomatitis virus (VSV) and lentiviruses were performed in a BSL-2+ setting following biosafety regulations of the Institut Pasteur, Paris. The VSV Indiana strain was kindly provided by N. Escriou (Institut Pasteur).

### TCID_50_ assays

Supernatants of infected cells were first cleared of cell debris by centrifugation at 500 rpm for 5 minutes at 4°C. They were 10-fold serially diluted in DMEM supplemented with 2% FBS and 1% P/S. Vero-E6 cells in suspension were infected with 10-fold serial dilutions in 96-well plates. Cells were seeded at a concentration of 80 000 cells/ml and incubated for 5 days in DMEM medium containing 2% FBS. At the end of the incubation period, cells were washed twice with PBS and then fixed with a crystal violet solution containing 3% formaldehyde for 30 minutes at room temperature. Cytopathic effects (CPE) were assessed by calculating the 50% tissue culture infective dose (TCID_50_) using the Spearman-Karber method [60].

### Cloning of host proteins and production of pseudoviruses

The sequence of *Rhinolphus malayanus* ACE2 (rACE2) was synthesized with a C-terminal V5-6 His sequence tag and was amplified using primers designed to add a 5’ BclI restriction site (rACE2 for: 5’-TCTAGTTGATCAGCCACCATGTCAGGCTCTAGCTGGC) and 3’ XhoI restriction site (rACE2-V5 rev: 5’-GAGAGGCTCGAGTCTATCAATGGTGATGGTG). Human ACE (hACE2) coding sequence was amplified from pLenti6-hACE2 previously described [61] with the same primer design strategy as for rACE2 (hACE2 for: 5’-TCTAGTTGATCAGCCACCATGTCAAGCTCTTCCTGGCTCCTTC, hACE2 rev: 5’-GAGAGGCTCGAGTCGAGTTAAAAGGAGGTCTGAACATCATCA). PCR products were digested using BclI/XhoI restriction enzymes and cloned in the pFlap-Ubc-nLuc-IRES-Puro lentiviral vector (kind gift from Pierre Charneau, Institut Pasteur, Paris, France) digested with BamHI/XhoI.

Human TMPRSS2 (hTMPRSS2) lentiviral vector was purchased from Addgene (pWPI-IRES-Bla-Ak-TMPRSS2; addgene plasmid #154982). *Rhinolophus ferrumequinum* TMPRSS2 (rTMPRSS2) coding sequence was amplified from *Rhinolophus ferrumequinum* cDNA using primers designed to add a N-terminal c-myc Tag and a 5’ PmeI restriction site (rTMPRSS2 fwd: 5’-TAGCCTCGAGGTTTAAACCCGGGAGCAGCACCATGGAGCAGAAACTCATCTCTGAAGAG GATCTG ATGGCTTTAAACTCAGGATC) as well as a 3’ SpeI restriction site (rTMPRSS2 rev: 5’-GTGGCCACTAGTACGTACGGTCCGCATATGGATCCCTAGCTGTTTGCCCTCATTTGTC).

Amplicon were cloned into the backbone of pWPI-IRES-Bla-Ak-TMPRSS2 lentiviral vector using PmeI/SpeI restriction sites. All plasmids were propagated in Stbl3 *E. coli* cells and verified by sequencing (Eurofins Genomics). To generate RFe-AT and RFe-ATC cells, plasmids pTRIP-SFFV-Hygro-2A-TMRPSS2 and pWPI-IRES-Bla-Ak-ACE2 Hygro_TMPRSS2 were used. The lentiviruses coding for the S protein of BANAL-236 and SARS-CoV-2 (BetaCoV/France strain) were previously described [1]. Lentiviral pseudovirions were produced in 293T cells in 10-cm dishes. Co-transfection included the spike plasmid (5 μg), a lentiviral vector expressing luciferase (10 μg, pHAGE-CMV-Luc2-IRES-ZsGreen-W), and auxiliary plasmids (3.3 μg each) HDM-Hgpm2, HDM-tat1b, pRC-CMV-Rev1b using calcium phosphate precipitation. After 5 hours, the medium was replaced with serum-free and phenol red-free DMEM. Pseudo-particles were harvested at 48 hours, clarified by centrifugation, and frozen. Control lentiviruses without S coding sequence were produced in parallel. The amount of pseudo-particles was quantified by p24 ELISA, as described previously [62].

### Cloning of BANAL-236 ORFs

BANAL-236 sequences were amplified from BANAL-236 cDNA using primers designed to add attb1 and attb2 recombination cassettes (Table S2). Difficult to amplify ORFs were synthetized flanked by attb1 and attb2 sequences. PCR products and synthetic DNA were cloned into pDONOR221entry vector using Gateway BP clonase II Enzyme Mix (Thermo Fisher Scientific). All entry clones were transferred into gateway compatible pciNeo 3X-Flag expression vector (kind gift of Yves Jacob, Institut Pasteur). BANAL-236 NSP3 coding sequence was amplified using primers designed to add 5’ NotI and 3’ BamHI sites (Table S2) and in-frame cloned into P3XFLAG-CMV-10 (Promega) using NotI /BamHI restriction enzymes. C-terminally STREP tagged BANAL-236 ORF8 and S expression vector were constructed cloning synthetic sequence (Table S2) into pLVX-EF1alpha-IRES-Puro vector (Takara) using EcoRI and BamHI restriction sites. Gene synthesis was performed by Genecust.

Alternatively, SARS-CoV2 and RaTG13 viral open reading frames (ORF) previously cloned in pDONR207 [28,63] were used as templates to derive BANAL-236 sequences using site directed mutagenesis (Quick-change site directed mutagenesis kit, Agilent Technologies Inc.) with primers listed in table S3. Previously cloned E, NSP9, NSP10 and NSP16 sequences of SARS-CoV2 in pDONOR207, which are identical to BANAL-236 (Table S1) were not modified.

### Spike-pseudotyped lentivirus entry assays

293T cells stably or transiently expressing hACE2, hTMPRSS2, or both entry factors, as well as RFe cells stably expressing hACE2, hTMPRSS2, rACE2, or a combination of entry factors, were transduced in suspension with pseudoviruses at 0.5 ng of HIV-1 p24 antigen for 2.10^4^ cells and then plated in 96-well plates. After 72h of transduction, luciferase activity was measured by adding 100 μl of Bright-Glo substrate (Promega) and quantifying luminescence with a Varioskan LUX luminometer (Thermo Fisher). The expression of the entry factors was assessed by flow cytometry at D0 for each experiment.

### Transfections

For poly I:C stimulation, RFe cells were plated in 12-well plates. The next day, they were transfected with 100 ng poly I:C (InvivoGen) or PBS, respectively, using INTERFERin (Polyplus Transfection) transfection reagent. Caco-2 cells were transfected with 500 ng of poly I:C using lipofectamine RNAiMAX (Thermo Fisher). Cells were lysed 16 h after transfection.

For siRNA transfection, cells were plated in 6-well plates and transfected the next day with 100 nM of siRNA targeting human TMPRSS2 (7113, Horizon Discovery Biosciences), using Lipofectamine RNAiMAX (Thermo Fisher) transfection reagent. Cells were harvested 72 h after transfection.

### Luciferase reporter assays

For luciferase experiments 293T cells were seeded in 96-well plates and transfected using Trans IT®-293 (Mirus) or using FuGENE® HD (Promega) with a mixture of 20 ng of pISRE-Luc, 2 ng of pRL-TK-Renilla and 20 ng of the plasmids expressing viral proteins or 20 ng of pCi-Neo plasmids (Promega) as “empty vector” (EV) controls. NS1 Flu (kindly provided by Daniel Marc, INRAE) and NS5 TBEV [42] were used as positives controls. All plasmids were grown in TOP10 cells (Thermo Fisher Scientific) and verified by sequencing. To stimulate the RIG-I/IRF3 axis, 20ng of pdeltaRIG-I (kindly provided by Pierre Genin, Centre d’Immunologie et des Maladies Infectieuses, Paris) was added in the plasmid mix. To activate the JAK/STAT pathway, cells were stimulated with 200 IU/ml of IFNα2b (PBL assay science) 8 hours post-transfection.Twenty-four hours post-transfection, cells were lysed using Passive Lysis buffer (Promega) for at least 15 min and luciferase activity was measured with Dual-Glo Luciferase Assay System (Promega) following the manufacturer’s protocol.

### Immunofluorescence microscopy

Cells were fixed with 4% paraformaldehyde (PFA) (Sigma-Aldrich) for 30 min at room temperature. Cells were blocked for 30 min with PBS containing 0.05% Saponin and 5% BSA before incubation with the indicated primary antibodies for 1 hour. After incubation, cells were washed three times with PBS containing 0.05% saponin and 5% BSA. Alexa Fluor 488 or 647 conjugated secondary antibodies were added for 1 hour. After incubation, cells were washed twice with PBS 0.05% saponin and 5% BSA and once with PBS. Nuclei were stained for 20 min with Hoechst-PBS (Thermo Fisher, 33342 (20 nM)). After washing, slides were mounted with Prolong Gold imaging medium (Life Technologies, P36930). Images were acquired using a Leica SP8 confocal microscope or Zeiss LSM 700 inverted.

### Flow cytometry

Cells were detached with trypsin and fixed in 4 % PFA for 30 min at 4°C. Staining was performed in PBS, 2 % bovine serum albumin (BSA), 2 mM EDTA (Invitrogen), and 0.1 % saponin (FACS buffer). Cells were washed tree time and incubated with indicated antibodies during 1 hour at 4°C. Cells were washed tree time again and incubated with secondary antibody Alexa 488 or 647 during 45 min in the dark at 4°C. For cell surface staining, cells were detached with trypsin and washed with PBS, 0.5 % BSA and 0.4 % EDTA (surface buffer). Cells were washed tree time and incubated with indicated antibodies (hACE2-Alexa 647, hTMPRSS2 [30] or anti-N [64]) during 1 hour at 4°C. For hTMPRSS2 staining, cells were washed tree time again and incubated with secondary antibody Alexa 647 during 45 min in the dark at 4°C. Cells were wash tree times again and fixed in 1 % PFA for 10 min at 4°C. Data were acquired on an Attune NxT flow cytometer (Thermo Fisher) and were analyzed using FlowJo software v10 (TriStar).

### Transmission Electron Microscopy

About 10^6^ Caco-2 cells were mock-infected or infected with SARS-CoV-2 or BANAL-236 at MOI 0.001 and 0.01 respectively. RFe cells were mock-infected or infected with SARS-CoV-2 or BANAL-236 at MOI of 0.02 and 0.2, respectively. Cells were detached with trypsin and then wash twice with PBS and incubated in 1% glutaraldehyde/4% paraformaldehyde (Sigma, St. Louis, MO) in 0.1 M phosphate buffer (pH 7.2). Samples were then washed in PBS and postfixed by incubation for 1 h with 2% osmium tetroxide (Agar Scientific, Stansted, UK). The cells were then fully dehydrated in a graded series of ethanol solutions and propylene oxide. They were impregnated with a 1:1 mixture of propylene oxide/Epon resin (Sigma) and left overnight in pure resin. Samples were then embedded in Epon resin (Sigma), which was allowed to polymerize for 48 h at 60°C. Ultrathin sections (90 nm) of these blocks were obtained with a Leica EM UC7 ultramicrotome (Wetzlar, Germany). Sections were stained with 2% uranyl acetate (Agar Scientific) and 5% lead citrate (Sigma), and observations were made with a transmission electron microscope (JEM-1011; JEOL, Tokyo, Japan).

### Western blot analysis

Cells were lysed in RIPA buffer (Sigma-Aldrich) supplemented with a protease and phosphatase inhibitor cocktail (Roche). Samples were denatured in 4X loading buffer (Li-Cor Bioscience) under reducing conditions (NuPAGE reducing agent, Thermo Fisher Scientific). Proteins were separated by SDS-PAGE (NuPAGE 4-12% Bis-Tris gel or NuPAGE 10-20% Tricine gel for small protein, Invitrogen) in running buffer MOPS (Invitrogen) or MES (Invitrogen) for small proteins. Proteins were transferred to nitrocellulose membranes (Bio-Rad) using a Trans-Blot Turbo system (Bio-Rad) or liquid transfer for NSP3. Membranes were blocked with PBS-Tween 0.1% containing 5% milk. After blocking, membranes were incubated overnight at 4°C with primary antibodies diluted in blocking buffer or PBS-Tween 0.1%. Finally, membranes were washed and incubated for 45 minutes at room temperature with diluted secondary antibodies, then washed three time again. Images were acquired with an Odyssey CLx infrared imaging system (Li-Cor Bioscience).

### Antibodies and cytokines

Details about antibodies are given in Table S4. Cells were stimulated overnight with IFNα2b (PBL Biosciences) at a final concentration of 1 000 IU/ml.

### RNA extractions and RT-qPCR assays

Total RNA was extracted from cell lysates using the NucleoSpin RNA II kit (Macherey-Nagel) and eluted in water. cDNA synthesis was performed on 1 µg of total RNA with RevertAid H Minus M-MuLV reverse transcriptase (Thermo Fisher Scientific) using random p(dN)6 primers (Roche). Real-time quantitative PCR was carried out on a Quant Studio 6 Flex system (Applied Biosystems) with SYBR green PCR mix (Life Technologies). Data were analyzed using the ΔΔCT method, with all samples normalized to GAPDH. All experiments were performed in technical triplicate. The primers used for RT-qPCR analysis are listed in Table S5. Genome equivalent concentrations were determined by extrapolation from a standard curve generated from serial dilutions of plasmids expressing hACE2, hTMPRSS2, rACE2-V5, C-Myc-rTMPRSS2 or BANAL-236 E protein.

### Data analysis

Data are presented and analyzed using GraphPad Prism 9. Alignments and trees were generated using CLC Genomics Workbench 22. Immunofluorescence quantification was done using Zen Blue software.

## Acknowledgments

We thank members of the Virus and Immunity Unit of the Institut Pasteur in Paris (Andréa Cottignies-Calamarte, Françoise Porrot, Florence Guivel-Benhassine, Julian Buchrieser, Nell Saunders, Mariem Znaidia and Nicoletta Casartelli) for reagents, advice, p24 titration and discussion. We also thank the French National Reference Centre for Respiratory Viruses hosted by Institut Pasteur (France) and headed at the time by Pr. S. van der Werf for providing the viruses.

---

**Figure S1.**
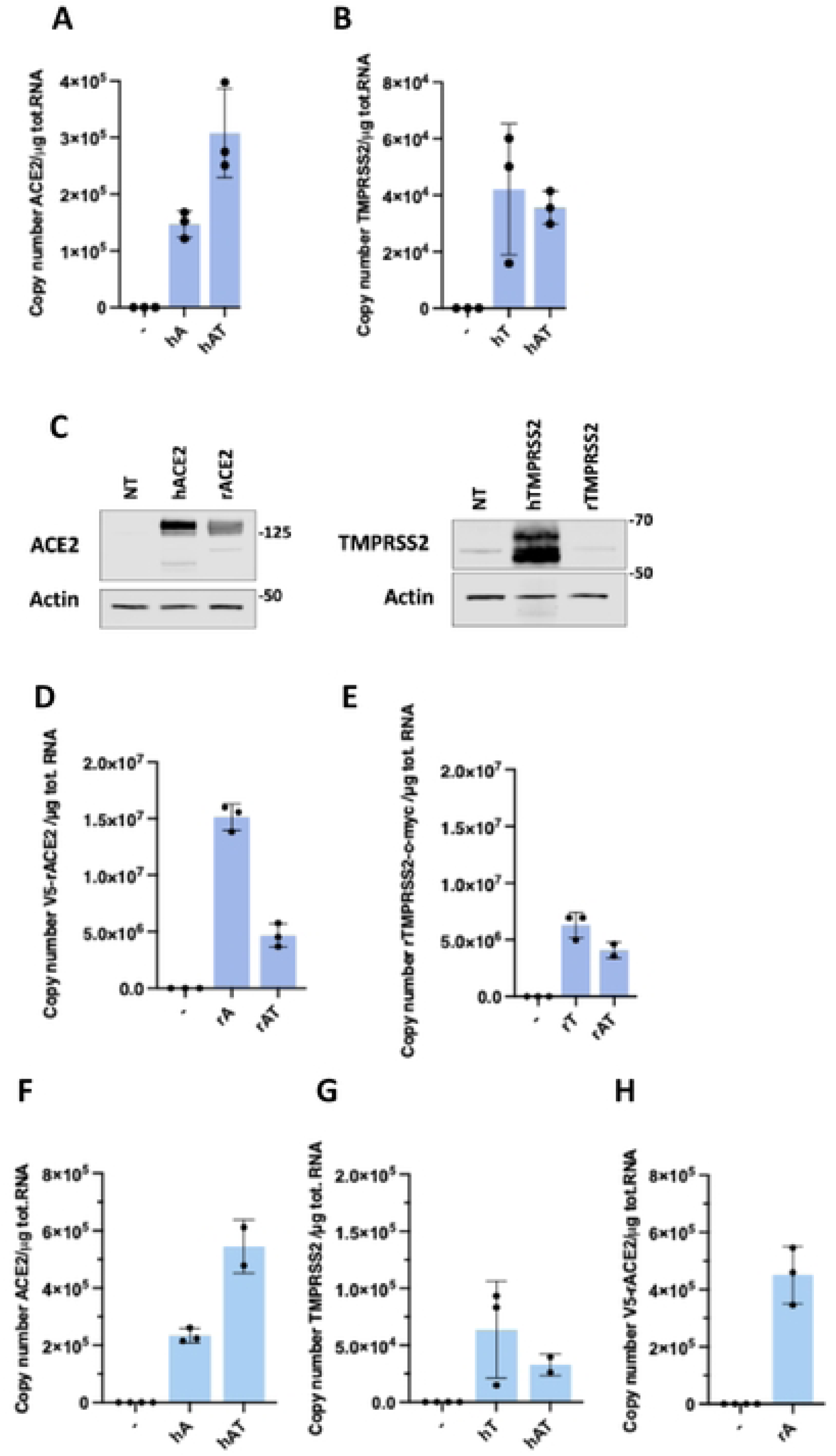
Expression of ectopically expressed ACE2 and TMPRSS2 in 293T and RFe cells. Quantification of copy numbers per µg of total cellular RNA of hACE2. (**A**) or hTMPRSS2 (**B**) in 293T cells via qPCR analysis. Data arc means ± SD of at least two independent experiments. (**C-D**) rACE2-V5 (**C**) or rTMPRSS-c-myc (**D**) in 293T cells. (**E-G**) RFc cells quantification of copy numbers per µg of total cellular RNA via qPCR analysis. Data for hACE2 (**E**), hTMPRSS2 (**F**) and rACE2-V5 (**G**). Data arc means ± SD of at least two independent experiments. (**H-I**) 293T cells were transfected for 24 h with plasmids expressing hACE2 or rACE2 (**H**) or expressing hTMPRSS2 or rTMPRSS2 (**I**). Whole-cell lysates were analyzed by western blotting with antibodies against the indicated proteins. Data are representative of three independent experiments. Quantification of copy numbers per µg of total cellular RNA of V5-rACE2.

**Figure S2.**
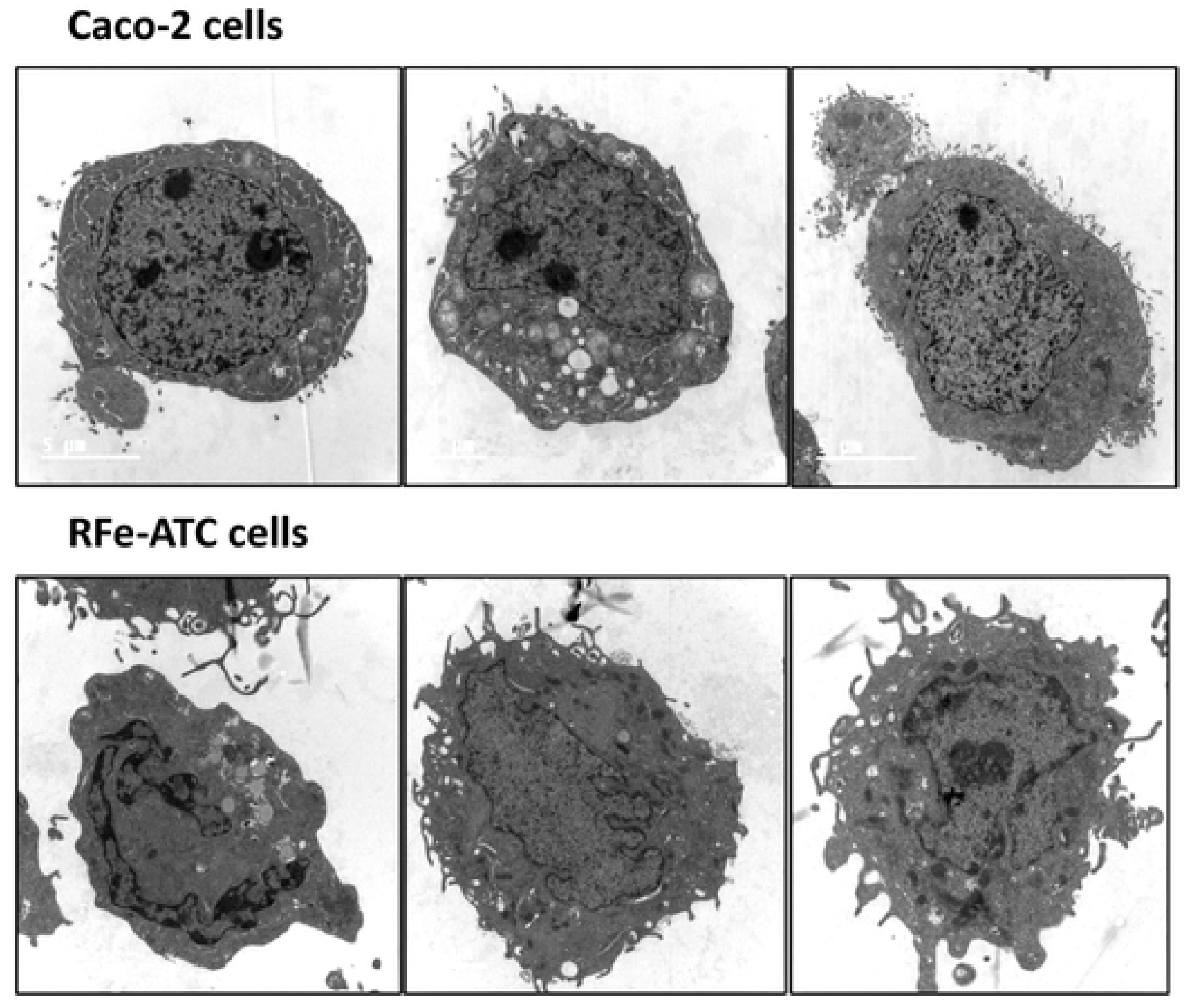
Transmission electron microscopy analysis of Caco-2 and RFe-ATC cells. Mock-infected cells were used as control cells for experiments shown in figure 5.

**Figure S3.**
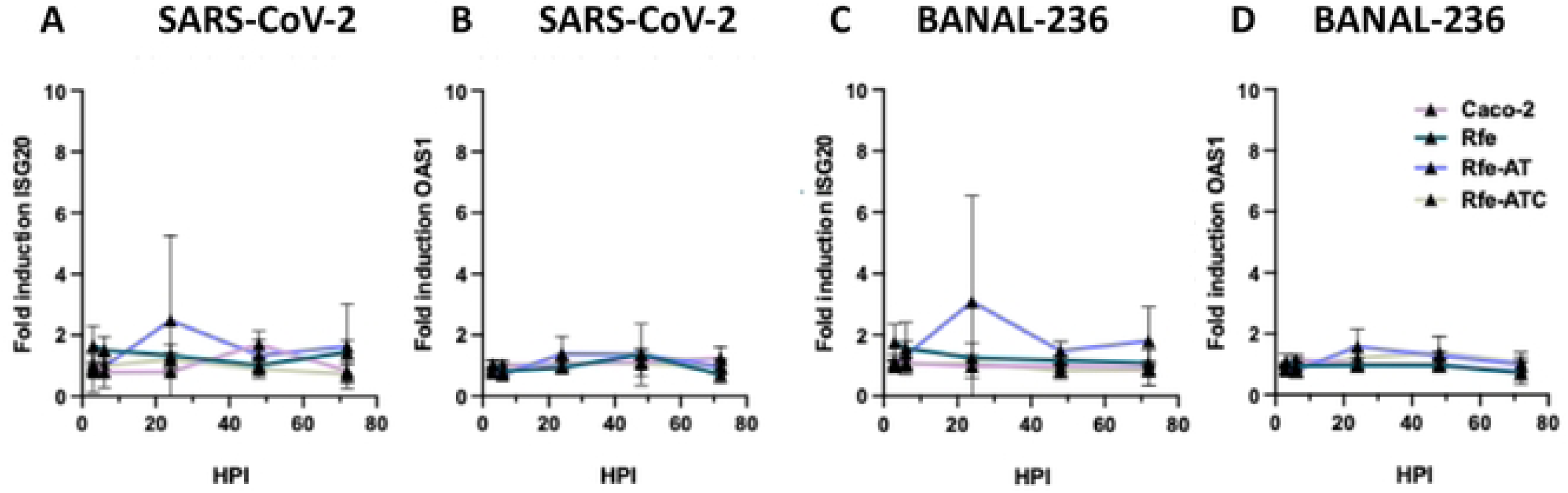
InteRFeron-stimulated genes are not induced upon SARS-CθV-2 and BANAL-236 replication in Caco-2 and RFe-ATC cells. RFe (green). RFc-AT (blue) and RFe-ATC (gray) cells were infected with SARS-CoV-2 at a multiplicity of infection (MOI) of 0.2 (**A, B**) or with BANAL-236 at an MOI of 0.5 (**C, D**). Caco-2 cells (pink), were infected with SARS-CoV-2 at an MOI of 0.0002 (**A, B**) or were infected with BANAL-236 at a MOI of 0.02 (**C, D**). Cell lysates were collected for RT-qPCR analysis at 3h. 6h, 24h. 48h, 72h. The relative amounts of 1SG20 (**A, C**) and OASI (**B, D**) mRNAs were determined by RT-qPCR analysis. Results were first normalized to GAPDH mRNA and then to mRNA levels of mock-infected cells, which were set at I. Data arc means ± SD of at least two independent experiments.

